# Peptide Sequence Determines the Efficacy of Single-Dose AAV-Vectorized GLP-1R Agonists in Diet-Induced Obese Mice

**DOI:** 10.64898/2026.11.08.759712

**Authors:** Donghyung Lee, Krister Bokvist, Greg Nachtrab, Adam Mezo, Tarsis Brust, Elijah Trefts, Julia Brusilovsky, Jiyoung Suk, Vincent Luczak, Lynn Edwards, Brooke Benedetto, Jude Onyia, David Woldbye

## Abstract

**Background:** Glucagon-like peptide-1 receptor (**GLP-1**R) agonists such as semaglutide produce substantial weight loss and glycemic improvement, but their long-term efficacy depends on sustained adherence to chronic dosing. Adeno-associated virus (AAV) gene therapy could, in principle, provide durable **GLP-1**R agonist (**GLP-1**RA) exposure from a single administration, but the degree of peptide engineering required for a vectorized GLP-1RA transgene to be efficacious in vivo is unknown.

**Methods:** We generated four self-complementary AAV (scAAV) constructs encoding: native human **GLP-1**(7-36) (**GLP-1**), a DPP-IV-resistant **Gly8**-**GLP-1**(7-36) variant (**Gly8**), Exendin-4 (Ex4), or an Exendin-4 human IgG4 Fc fusion protein (Ex4-Fc). All four constructs were validated to secrete functional **GLP-1**R agonists in vitro using a conditioned-media potency assay. Subsequently, these constructs were packaged into PHP.eB capsids and administered as a single intravenous dose (5×10^12^ vg/kg) to diet-induced obese (DIO) mice alongside a vehicle control. Body weight, body composition, food intake, fasting plasma insulin, and oral glucose tolerance were assessed over a 3-week period. At the end of the study, tissues were harvested for assessments of plasma/liver lipids, tissue transgene transcript and peptide levels.

**Results:** HEK293 cells transfected with the four AAV constructs all secreted active peptides/proteins as assessed by a cell-based **GLP-1**R activation assay, although the effective concentration of the **GLP-1** constructs in the media spanned nearly four orders of magnitude 72 hours after transfection. In diet-induced obese (DIO) mice, **Ex4** and **Ex4-Fc** vectors, but not **GLP-1** or **Gly8** vectors, produced significant reductions in body weight of up to 22.6% of starting body weight out to 21 days after a single dose. In addition, reductions in food intake, fat mass, and fasting insulin relative to vehicle were also observed for **Ex4** and Ex4-Fc. Furthermore, **Ex4-Fc** improved oral glucose tolerance while reducing plasma cholesterol and liver triglycerides relative to vehicle. Although transcript levels did not differ appreciably among the four constructs in either brain or liver, measurable peptide product in plasma and brain lysate tracked the rank order of the in vitro supernatant screen, with **GLP-1** and **Gly8** groups indistinguishable from endogenous background.

**Conclusions:** PHP.eB scAAV **Ex4** and **Ex4-Fc** showed robust efficacy on multiple metabolic endpoints including weight loss and glucose tolerance. In contrast, vectorized **GLP-1** and **Gly8** showed no in vivo efficacy despite comparable levels of viral transcript in the brain and the liver. Vectorized **GLP-1** and **Gly8** also performed poorly compared to exendin-4-based constructs in vitro, suggesting that conditioned-media potency screening is a valid early filter for construct selection in vectorized-peptide AAV development. Engineering peptides to improve translation and secretion efficiency, in addition to circulating plasma stability, could further optimize vectorized **GLP-1**RA therapeutics.

## Introduction

Obesity and type 2 diabetes are leading drivers of cardiometabolic morbidity worldwide. Pharmacologic **GLP-1** receptor agonism is now the most effective non-surgical treatment available for obesity: once-weekly semaglutide produces mean weight loss approaching 15% over 68 weeks [1], and the dual glucose-dependent insulinotropic polypeptide (GIP)/**GLP-1** receptor agonist tirzepatide produces even larger reductions [2]. The shift from earlier twice-daily formulations to once-weekly dosing has meaningfully reduced the treatment burden of **GLP-1**R agonist (**GLP-1**RA) therapy, but efficacy still depends on indefinite continuation of treatment. Roughly two-thirds of lost weight, and the accompanying cardiometabolic benefit, is reversed within a year of stopping the treatment [3,4]. Because a portion of the weight lost during **GLP-1**RA therapy is lean mass [5], repeated cycles of treatment and discontinuation have raised concern for sarcopenic obesity [6]. Furthermore, real-world studies report high discontinuation rates for semaglutide and tirzepatide [7,8]. These limitations bolster the case for a single-administration vectorized therapeutic that produces durable **GLP-1**R engagement and lifelong clinical efficacy without chronic dosing.

Native **GLP-1**(7-36) is a poor candidate for vectorized **GLP-1**RA therapy because it is cleaved by dipeptidyl peptidase-IV (DPP-IV) at its N-terminal Ala8-Glu9 bond and by neutral endopeptidase 24.11 (NEP) within minutes of entering the circulation, resulting in a plasma half-life of only a few minutes [9,10]. The clinically approved **GLP-1**RAs circumvent this liability in two ways: with amino acid substitutions at position 8 to confer DPP-IV resistance and with fatty-acid conjugation to confer binding to serum albumin and subsequent half-life extension [11]. Liraglutide utilizes fatty-acid conjugation whereas semaglutide and tirzepatide incorporate the unnatural amino acid Aib at position 8 as well as fatty-acid conjugation. In contrast, exendin-4-based therapeutics (exenatide) instead use a *Heloderma suspectum*-derived sequence that is intrinsically resistant to both DPP-IV and NEP [12,13]. Both strategies pose problems in a gene therapy context: neither fatty-acid acylation nor unnatural amino acids can be installed by the ribosome and sequence-level engineering introduces the risk of anti-drug antibody (ADA) formation and subsequent neutralization, clearance of the transgene product, or immune clearance of transduced cells [14]. A vectorized **GLP-1**RA built from the native **GLP-1** would minimize ADA risk, but it remains unclear whether constitutive local production of native **GLP-1** could overcome its known in vivo instability.

We characterized the degree to which engineering affects vectorized GLP-1RA efficacy by designing and expressing a series of GLP-1RA sequences, differing in their resistance to peptidase degradation and systemic clearance, from an otherwise identical AAV expression cassette. We used the PHP.eB capsid because it transduces both the central nervous system and peripheral organs, including liver, after intravenous administration in C57BL/6J mice [15,16]. This allowed for the evaluation of the transgene product in compartments relevant to both central (hypothalamic and hindbrain) and peripheral (pancreatic beta-cell, vagal afferent) mechanisms of GLP-1RA action [17,18,19,20].

To this end, we compared four AAV-delivered GLP-1RAs in diet-induced obese mice: 1) native human **GLP-1**(7-36) (**GLP-1**); 2) **Gly8**-**GLP-1**(7-36) (**Gly8**), which carries a single alanine-to-glycine substitution at position 8 (A8G) that confers DPP-IV resistance without addressing NEP-mediated clearance [21]; 3) Exendin-4 (Ex4), which is resistant to both proteases; and 4) an Ex4–human IgG4 Fc fusion (Ex4-Fc), in which an antibody Fc domain is added to extend systemic half-life beyond the renal filtration threshold as has been demonstrated for dulaglutide [22]. By holding the expression cassette constant and varying only the GLP-1RA sequence, we sought to characterize how modifications in the GLP-1RA sequence itself impact efficacy in the context of continuous local production, and to relate any differences in efficacy to measured differences in tissue and circulating transgene product.

## Materials and Methods

### Ethics Statement

All procedures were conducted under an institutionally approved Institutional Animal Care and Use Committee (IACUC) protocol at Neurocrine Biosciences, Inc. Reporting follows the ARRIVE 2.0 guidelines to the extent the underlying study records allow.

### Vector Construct Design

Four expression cassettes encoding vectorized GLP-1RAswere designed for packaging into self-complementary AAV (scAAV) vectors. Each cassette was flanked by AAV2 inverted terminal repeats (ITRs), with the 5⍰ ITR containing a deleted terminal resolution site (ΔTRS) to enable self-complementary genome packaging [23]. Transgene expression was driven by the cytomegalovirus (CMV) immediate-early promoter/enhancer [24]. All constructs included the murine proglucagon signal peptide upstream of the GLP-1RA coding sequence to direct secretion. The 3⍰ untranslated region contained either the full-length woodchuck hepatitis virus posttranscriptional regulatory element (WPRE) [25] or a truncated variant (WPRE3) [26], followed by the bovine growth hormone polyadenylation signal (bGH polyA) [27]. Construct architecture and the GLP-1RA sequence alignment are shown in Figure 1A–B.

**Figure 1.**
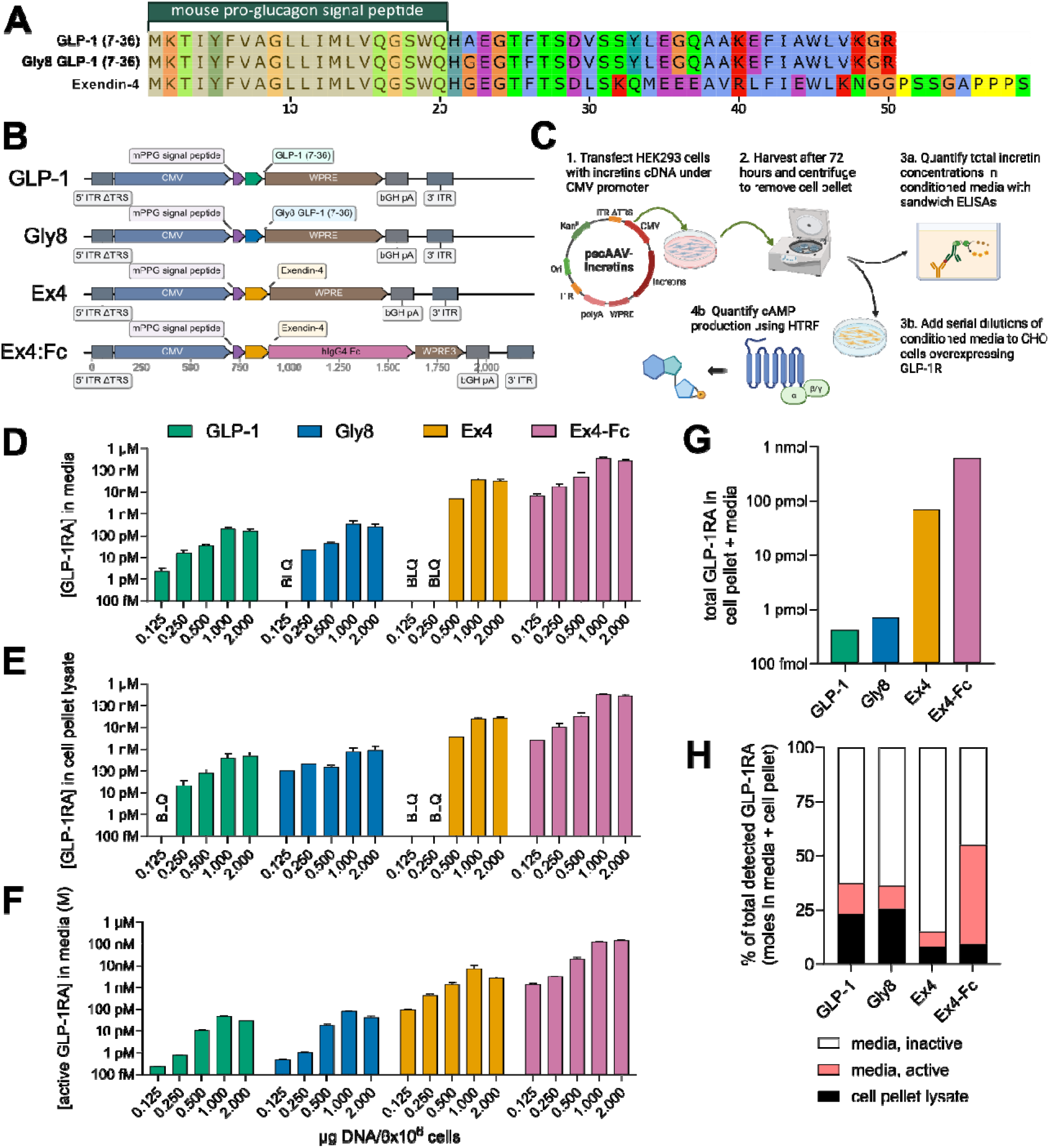
Vector construct design and in vitro validation. (A) Alignment of the full-length (signal peptide + mature GLP-1RA) **GLP-1, Gly8**, and **Ex4** sequences (**Ex4-Fc** contains the Exendin-4 sequence shown, fused via a linker to a human IgG4 Fc domain not depicted in the alignment). (B) Schematic architecture of the four AAV expression cassettes (5⍰ ITR to 3⍰ ITR). (C) In vitro workflow. (D) GLP-1RA concentration in media by ELISA 72 hours after transfection protocol. (E) GLP-1RA concentration in lysate by ELISA 72 hours after transfection protocol. (F) Receptor active GLP-1RA concentration in media (cAMP-interpolated). Bars show average of technical duplicates ± SEM, BLOQ = below limit of quantification. (G) Total GLP-1RA produced per construct at the 2 µg DNA dose, calculated as moles detected in cell pellet lysate plus moles detected in media (ELISA concentration x volume for each). (H) Partitioning of total GLP-1RA from G into three fractions (% of total) at the 2 µg DNA dose: cell pellet lysate, active media (from the **GLP-1**R functional assay), and inactive media (the ELISA − functional assay difference).

Coding sequences for the murine proglucagon signal peptide and human **GLP-1**(7-36) used endogenous mouse cDNA sequences. The **Gly8** construct was generated by substituting the second codon of the mature **GLP-1**(7-36) sequence with GGA. The **Ex4** coding sequence was reverse-translated from the peptide sequence using Benchling’s codon optimization tool. Signal-peptide cleavage between the signal-peptide glutamine and the mature GLP-1RA N-terminal histidine was confirmed computationally for all four constructs using SignalP 6.0 [28].

### Molecular Cloning and Plasmid Production

Gene fragments were synthesized by Twist Bioscience and assembled into the scAAV backbone by Gibson assembly using NEB HiFi DNA Assembly Master Mix (New England Biolabs). Final plasmid sequences were verified by Oxford Nanopore long-read whole-plasmid sequencing (Plasmidsaurus). ITR-to-ITR cassette sequences are available on reasonable request.

### Transient Expression and Peptide Quantification In Vitro

Viral Production Cells 2.0 (VPC2.0; Thermo Fisher Scientific, Gibco A49784), a clonal suspension-adapted HEK293F-derived line, were maintained in Viral Production Medium (Thermo Fisher Scientific) supplemented with 4 mM GlutaMAX (Thermo Fisher Scientific, 35050038) and 100 U/mL penicillin–streptomycin. To compare relative expression and bioactivity of each construct, VPC2.0 cells were seeded at 3×10^6^ cells/mL (2 mL/well, 24-well deep-well plates) and transfected with FectoVIR (Sartorius; 2 µL/well) at plasmid DNA inputs of 0, 0.125, 0.25, 0.5, 1.0, and 2.0 µg/well. All four constructs were transfected on the same plate to control for inter-assay variability. Cells were incubated for 72 h without media exchange (37°C, 8% CO2, 80% humidity, 650 rpm orbital shaking).

Cells were harvested by centrifugation (300 x g, 10 min, 4°C). The supernatant was supplemented with 1x Halt Protease Inhibitor Cocktail (Thermo Fisher, 78429) then 0.2 µm filtered to produce cell-free media for ELISA and cAMP assays. The cell pellets were lysed in 200 µL NP-40 lysis buffer (Thermo Fisher, J62805.EQE) with 1x Halt Protease Inhibitor then clarified by centrifugation (12,000 x g, 10 min, 4°C).

### ELISA Quantification

GLP-1RA concentrations in supernatant and lysate were measured by ELISA. **Ex4** and **Ex4-Fc** were measured with an Exendin-4 ELISA kit (Abcam, ab272192) at 1:100–1:4000 dilutions. **GLP-1** was measured with a Total **GLP-1** ELISA kit (MilliporeSigma, EZGLP1T-36K; detects both 7-36 and 9-36 forms) at neat and 1:3 dilution. Because this kit’s standards are not validated for the **Gly8** substitution, **Gly8** was instead interpolated from a standard curve of an albiglutide fragment (peptide portion of the A8G substituted **GLP-1** analog), run in parallel with the manufacturer standards. Samples were run in duplicate; concentrations were interpolated from a four-parameter logistic (4PL) fit (GraphPad Prism) and dilution-corrected. **Ex4-Fc** values were divided by 2 to account for Fc-mediated dimerization. Absorbance was read on a PHERAstar FSX (BMG Labtech).

### GLP-1R Activation Assay

Biological activity of secreted GLP-1RAs was assessed using the LANCE Ultra cAMP assay (PerkinElmer, TRF0262) in CHO cells stably expressing human **GLP-1**R, maintained in Ham’s F-12 (Gibco, 11765-054) with 10% FBS, 50 U/mL penicillin–streptomycin, and 400 µg/mL G418. Cells were resuspended in stimulation buffer (Hanks’ balanced salt solution, 4.5 mM HEPES, 0.45 mM IBMX, BSA) and seeded at 3,500 cells/well (384-well, Corning CLS3572). Supernatants were tested neat and at 1:20, 1:400, and 1:8,000. Reference standards (**GLP-1**(7-36) amide acetate, MedChemExpress HY-P0054; albiglutide peptide fragment, HY-108795; Exendin-4 acetate, HY-13443A; and in-house Protein A/G-purified **Ex4-Fc**) were run as 15-point, 3-fold serial dilutions from 10 nM. Cells were stimulated for 30 min at room temperature; time-resolved fluorescence was read on a PHERAstar FSX (337/620/665 nm TR-FRET module, 60–500 µs integration). Signals (the amount of cAMP induced relative to the response produced by known concentrations of each peptide standard) were converted to an apparent “functional” (**GLP-1**R-active) GLP-1RA concentration by 4PL interpolation against these peptide-specific standard curves; the most dilute sample within the quantifiable range was used for each determination. All samples were run in technical duplicate.

### AAV Production and Purification

Recombinant scAAV vectors were produced by triple transfection of VPC2.0 cells (Gibco, A49784) as previously described [29]. Cells (3×10^6^ cells/mL, 80 mL in 250-mL shake flasks) were transfected with PHP.eB capsid plasmid, pALD-HELP (Aldevron, 5082-10), and the respective transfer plasmid (0.14:0.43:0.43 molar ratio; 1 µg total DNA/10^6^ cells) using FectoVIR (Sartorius). All four constructs were processed identically and in parallel to limit batch-to-batch variability.

At 72 h post-transfection, cultures were lysed with AAV-MAX Lysis Buffer (Gibco, A50520) plus Benzonase (Merck, 104445; 100 U/mL) and 2 mM MgCl2 (37°C, 1.5 h). Lysates were clarified (3,000 x g, 15 min, 4°C) and precipitated overnight with 25% v/v 40% PEG 8000/2.5 M NaCl, then pelleted (3,000 x g, 30 min, 4°C) and resuspended in 1x gradient buffer.

AAV was purified by iodixanol density-gradient ultracentrifugation adapted from Zolotukhin et al. [30]. Discontinuous step gradients (15/25/40/58% iodixanol) were centrifuged at 63,000 rpm for 1 h at 18°C (Type 70 Ti rotor, Beckman Coulter, Optima XPN-100). The 40%/58% interface fraction was collected by side puncture and buffer-exchanged into formulation buffer (20 mM Tris-HCl pH 8.0, 192 mM NaCl, 2 mM MgCl2, 0.005% Poloxamer 188) using 100-kDa MWCO Amicon Ultra-15 filters to a final volume of ∼100 µL. Aliquots were reserved for titer determination and mass photometry; all material was stored at −80°C.

### AAV Genome Titer Determination by Digital PCR

Encapsidated genome titers were determined by digital PCR (QIAcuity Eight, Qiagen; 8.5k Nanoplate cartridges), adapted from Lock et al. [31]. Samples were treated with DNase I (NEB, M0303L) to degrade unencapsidated DNA, then Proteinase K-digested and heat-inactivated. Genomes were serially diluted and quantified with the WPRE/WPRE3-shared primer/probe set described above, using QIAcuity Probe PCR Master Mix (Qiagen, 250103), with HpaII restriction enzyme (ThermoFisher, ER0512) included to linearize self-complementary genomes prior to amplification. The dilution with 95% CI < 5% was used for titer calculation (vg/mL).

### Mass Photometry

AAV preparations were characterized by mass photometry to estimate the ratio of empty to full capsids [32], using a SamuxMP mass photometer (Refeyn) with MassGlass slides (MP-CON-21022), 1:10 dilution in formulation buffer, 60-s acquisition, and Refeyn DiscoverMP analysis. Particle populations were assigned as empty or full by comparison to an empty-capsid reference standard and expected molecular masses calculated from genome size (empty PHP.eB capsid: ∼3,750 kDa; estimated full-capsid masses: **GLP-1/Gly8**, ∼4,909 kDa; **Ex4**, ∼4,927 kDa; **Ex4-Fc**, ∼5,180 kDa). Mass photometry indicated similar full-capsid content across preparations (%full: 90.7, 92.2, 85.9, and 92.2 for **GLP-1, Gly8, Ex4**, and **Ex4-Fc**, respectively).

### Animals and Housing

Male diet-induced obese (DIO) C57BL/6NTac mice (Taconic Biosciences) were maintained on 60% kcal-from-fat high-fat diet (Research Diets D12492) from 6 weeks of age [33,34]. Mice arrived at 20 weeks of age, were single-housed (12:12 h light/dark, lights on 0600 h) with ad libitum HFD and water and habituated for 2 weeks before study initiation.

### Study Design, Randomization, and Sample Size

At 21 weeks of age, baseline oral glucose tolerance test (OGTT), body composition (EchoMRI-900) [35], body weight, and food intake were collected. Mice were randomized into five groups using the *balanced_study* Stratified Clustering Hybrid algorithm [36]: baseline metrics (OGTT area under the curve (AUC), body-fat percentage, body weight, and food intake) were z-scored, decorrelated by principal component analysis, clustered in principal-component space by k-means++, and distributed across groups by simulated annealing to jointly minimize between-group mean differences, within-group variance, and between-group Mahalanobis distance; group equivalence at baseline was confirmed by one-way ANOVA/Kruskal–Wallis per metric (Bonferroni-corrected) and by MANOVA. Group sizes at randomization were n = 7 (**GLP-1, Gly8, Ex4-Fc**, vehicle) and n = 6 (**Ex4**); after post hoc exclusion of animals with undetectable transcript in both tissues, n = 5–7 per group were analyzed.

### AAV Administration

At 22 weeks of age, mice received a single intravenous tail-vein injection of AAV diluted to 3×10^12^ vg/mL in formulation buffer, dosed at 5×10^12^ vg/kg (72–88 µL per mouse by individual body weight). Vehicle animals received an equivalent volume of formulation buffer.

### In-Life Assessments

Body weight and food intake were measured daily for week 1, then weekly through week 3; food intake was determined per animal from hopper weight loss. OGTTs were performed at baseline and 3 weeks post-dose: following a 5-h fast, mice received 20% oral glucose (Teknova, G0530; 2 g/kg), with blood glucose measured at 0, 15, 30, 60, 90, and 120 min (AccuChek Guide Me, Roche) [37]. Body composition (fat/lean mass) was assessed by quantitative NMR using an EchoMRI-900 (EchoMRI™) a week before AAV administration and 3 weeks post-dose [35]. Investigators were not blinded to group allocation during in-life assessments; body weight, food intake, body composition and blood glucose were recorded by instrument readout rather than subjective scoring.

### Blood Collection, Tissue Collection, and Homogenization

Interim fasted plasma (1 week pre-dose and 3 weeks post-dose; 5-h fast) was collected by tail snip into EDTA Microvette tubes (Sarstedt). Terminal blood was collected from ad libitum-fed mice by cardiac puncture under isoflurane anesthesia into EDTA K3E tubes. Mice were then transcardially perfused with saline to clear residual blood before liver and whole-hemisphere brains were dissected, snap-frozen, and stored at −80°C. Tissue was homogenized in PBS with 1x Halt Protease Inhibitor Cocktail using a Precellys Evolution homogenizer (Bertin Instruments); the same homogenate was used for protein, ELISA, and RNA analyses.

### Biochemical and Molecular Analyses

Fasted plasma insulin was measured by Mouse Ultrasensitive Insulin ELISA (ALPCO, 80INSMSU-E01). Plasma and liver total cholesterol and triglycerides were measured with Infinity reagent kits (Thermo Fisher, TR13421/TR22421); liver lipids were normalized to total protein (Pierce BCA, Thermo Fisher, 23225). Tissue and plasma **Ex4** (Abcam, ab272192) and Total **GLP-1** (Millipore Sigma, EZGLP1T) were measured by ELISA as above; brain values were protein-normalized and plasma values reported as dilution-corrected molar concentrations. Total RNA was extracted (Maxwell RSC simplyRNA Tissue Kit, Promega, AS1340) and quantified (Qubit Broad-Range RNA Assay, Invitrogen). Viral transgene transcript was measured by QIAcuity OneStep RT-dPCR targeting the shared WPRE/WPRE3 region (same primer/probe set as the genomic titer assay; ∼1 ng total RNA/reaction), reported as copies/ng RNA. Animals with undetectable transcript levels in both tissues were excluded post hoc from all analyses (at most one per group).

## Statistical Analysis

Data were analyzed in GraphPad Prism. One-way ANOVA was used for single-endpoint comparisons across the five groups; two-way (repeated measures where applicable) ANOVA was used for time-course data (body weight, food intake, OGTT). Post hoc comparisons to vehicle used Dunnett’s test unless otherwise noted in the Results. Significance was defined as p < 0.05. Exact group sizes for each analysis are given in the corresponding figure.

## Results

### Incremental Engineering of Vectorized GLP-1RAs

We designed four scAAV expression cassettes that were identical apart from the GLP-1RA coding sequence (Figure 1A): 1) **GLP-1**; 2) **Gly8**, carrying a single A8G substitution known to confer DPP-IV resistance while retaining **GLP-1** activity; 3) **Ex4**, intrinsically resistant to both DPP-IV and NEP [12,13] and 4) **Ex4-Fc**, in which exendin-4 was linked to the human IgG4 Fc domain via a mouse codon-optimized (G4S)x3 amino acid linker and S228P [38], L235E [39] mutations in the Fc region (Figure 1B). All constructs used the same CMV promoter, murine proglucagon signal peptide, and bGH polyA. However, since the **Ex4-Fc** cassette exceeded packaging constraints with the full-length WPRE, it used the shorter WPRE3 element [26], while the other three constructs retained full-length WPRE [25]. The RT-dPCR primer/probe set used for transcript quantification targets a sequence common to both WPRE and WPRE3, allowing the same assay to be used across all four constructs. ITR-inclusive cassette sizes were 1,829 bp (**GLP-1** and **Gly8**), 1,856 bp (**Ex4**), and 2,239 bp (**Ex4-Fc**).

### In Vitro Secretion and Activity on GLP-1R

After plasmid production, we assessed each vector’s ability to transduce peptide/protein production and functional secretion in vitro. To this end, we transfected VPC2.0 cells with varying amounts of DNA then collected media and cell lysates 72 h post-transfection for ELISA and **GLP-1**R cAMP assays (workflow, Figure 1C). All four constructs showed dose-related increases in total-GLP-1RA concentrations in both supernatant and lysate, suggesting that all constructs can support secretion of the intended peptides/proteins (Figure 1D–E). At the highest DNA input tested (2.0 µg), total concentrations in media were 100-to 1,000-fold higher for the exendin-4-based constructs compared to the **GLP-1**-based constructs (0.160 nM, 0.266 nM, 32.3 nM, 282 nM for **GLP-1, Gly8**, Ex4, and Ex4-Fc, respectively, Figure 1D). We note that the media concentration of **Ex4-Fc** at the lowest DNA input (0.125 µg) exceeded media concentration of **GLP-1** at the highest DNA input (2.0 µg) (6.6 nM vs 477 pM, respectively). The GLP-1RA levels measured in cell lysates at 2.0 µg DNA input closely matched the pattern observed in media (0.477 nM, 0.911 nM, 27.5 nM, 278 nM for **GLP-1, Gly8**, Ex4, and Ex4-Fc, respectively, Figure 1E).

The immunoassays used to quantify peptide/protein concentrations cannot distinguish between active and inactive GLP-1RAs. To assess whether the GLP-1RAswere functionally active in the conditioned media, we tested the conditioned media in a cAMP accumulation assay using **GLP-1**R overexpressing CHO cells. Media from all four constructs produced dose-related **GLP-1**R-mediated cAMP responses in CHO-**GLP-1**R cells, validating these constructs’ abilities to generate functionally active GLP-1RAs(Figure 1F). By comparing the cAMP responses to a standard curve, we interpolated the functional GLP-1RA concentrations at the highest DNA input (2.0 µg) to be 29.6 pM, 42.6 pM, 2.73 nM, 147.5 nM for **GLP-1, Gly8**, Ex4, and Ex4-Fc, respectively, matching the total-concentration rank-order measured by the immunoassays.

At the highest DNA input (2.0 µg), total GLP-1RA production (lysate + media combined) varied more than 1,000-fold across constructs, from 414 fmol for **GLP-1** to 620 pmol for **Ex4-Fc** (Figure 1G). Partitioning this total into cell pellet lysate, **GLP-1**R active-media, and inactive-media fractions revealed differences in secretion and bioactivity across the four constructs (Figure 1H). The exendin-4-based constructs retained a smaller fraction of total product in the cell pellet lysate (7.85% and 8.99% for **Ex4** and Ex4-Fc, respectively) than **GLP-1** (23.0%) and **Gly8** (25.5%), indicative of a 2-to 3-fold improvement in secretion efficiency. **Ex4-Fc** also had the largest active-media fraction (45.8% of total), suggesting up to 6.5-fold better protein-level stability under these in vitro conditions. We note, however, that these differences in secretion efficiency and bioactivity are orders of magnitude smaller than the ∼1,000-fold differences in overall production described above.

Taken together, in vitro characterization identified **Ex4-Fc** as the most efficiently produced construct. In addition to total protein production up to ∼1,000-fold higher than the worst-performing **GLP-1** construct, **Ex4-Fc** had the largest fraction of active GLP-1RA in media and the smallest fraction of total product retained intracellularly (Figure 1G–H). Despite these differences, all four constructs were validated to secrete functional GLP-1RAsin vitro, supporting further evaluation of all four constructs in vivo.

### AAV Vector Production, Body Weight, Body Composition, and Food Intake

GLP-1 is known to have both central and peripheral mechanisms of action. To ensure that we transduce cells in both compartments, we chose to package all four constructs into the PHP.eB capsid, which can transduce both the central nervous system and peripheral organs after intravenous administration in C57BL/6J mice [16] (Figure 2A). Digital-PCR titers of the four preparations (200 µL final formulated volume from a 30 mL production culture) were 1.77×10^13^, 1.17×10^13^, 7.41×10^12^, and 1.48×10^13^ vg/mL for **GLP-1, Gly8**, Ex4, and Ex4-Fc, respectively. All vectors were administered at a nominal dose of 5×10^12^ vg/kg, selected based on prior characterization of the PHP.eB capsid, in which a comparable dose transduced the majority of cortical and striatal neurons following systemic administration [16].

**Figure 2.**
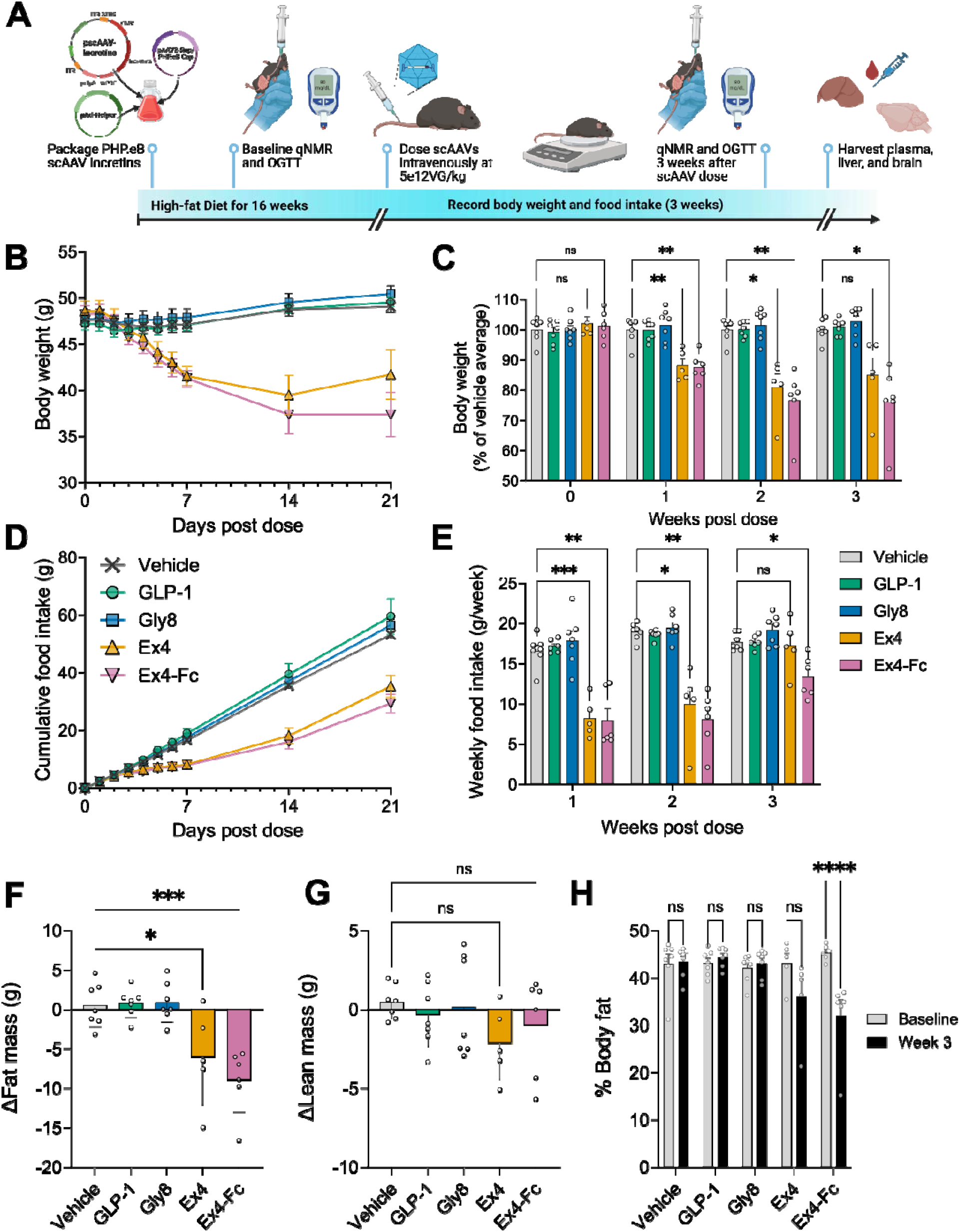
Body weight, body composition, and food intake after a single intravenous dose of vectorized GLP-1RA AAV in diet-induced obese mice. (A) In vivo workflow. (B) Body weight (g) over 21 days post-dose. (C) Weekly body weight (% of vehicle average) across the 3-week study (D) Cumulative food intake across 21 days post-dose. (E) Weekly food intake, weeks 1–3. (F) Change in fat mass (g) from baseline to week 3 (qNMR). (G) Change in lean mass (g) from baseline to week 3. (H) Percent body fat at baseline versus week 3. Data were analyzed by two-way repeated-measures ANOVA with Dunnett’s post hoc tests versus vehicle at each time point (C, E), one-way ANOVA with Dunnett’s post hoc tests versus vehicle (F, G), or two-way repeated-measures ANOVA with Šídák’s post hoc tests comparing baseline versus week 3 within each group (H) n = 5–7/group; ns, not significant; *p < 0.05; **p < 0.01; ***p < 0.001; ****p < 0.0001.

Vectors were administered intravenously to 22-week-old male DIO mice (Figure 2A). Body weights were similar across all groups before dosing and up to 2 days after dosing. By day 3, animals dosed with the **Ex4** and **Ex4-Fc** vectors began to diverge in body weight from the vehicle group. By day 7, **Ex4** and **Ex4-Fc** groups had respectively lost an average of 14.5% and 14.4% of starting body weight, while the **GLP-1** and **Gly8** groups’ weights were maintained, much like the vehicle group’s (Figure 2B). Body weight in both exendin-4-based groups remained significantly reduced until the 3-week study endpoint, reaching 85.0% and 76.1% relative to vehicle for **Ex4** and Ex4-Fc, respectively (p < 0.001 and p < 0.0001, Figure 2C). Between weeks 2 and 3, average body weight in the **Ex4** group increased modestly from a week-2 nadir (from 81.2% to 85.7% of starting body weight), whereas the **Ex4-Fc** group remained stable over the same interval (Figure 2C).

Consistent with the weight findings, cumulative food intake was reduced in the **Ex4** and **Ex4-Fc** groups relative to vehicle (Figure 2D). On a weekly basis, food intake was significantly reduced relative to vehicle in both groups during weeks 1 and 2 (Ex4: p < 0.001, p < 0.05; Ex4-Fc: p < 0.01, p < 0.01) (Figure 2E). By week 3, food intake in the **Ex4** group was no longer significantly different from vehicle, correlating with the week-3 rebound seen in body weight. The **Ex4-Fc** group’s food intake also increased from week 2 to week 3 but remained significantly reduced relative to vehicle (p < 0.05). **GLP-1** and **Gly8** groups did not differ from vehicle in food intake at any time point.

To further describe the nature of weight loss, we also measured lean mass and fat mass across this 3-week period using EchoMRI. Loss of fat mass from baseline was significantly higher relative to vehicle in both the **Ex4** (p < 0.01) and **Ex4-Fc** (p < 0.0001) groups (Figure 2F). The change in lean mass did not differ significantly between any treatment group and vehicle, though **Ex4** and **Ex4-Fc** groups trended lower than vehicle and **GLP-1**/**Gly8** groups (Figure 2G). When body fat percentage at week 3 was compared to each group’s own baseline, only the **Ex4-Fc** group reached statistical significance (baseline ∼45% to week 3 ∼32%, p < 0.0001) (Figure 2H). The change in body fat percentage for the **Ex4** group did not reach statistical significance, despite a similar downward trend.

### Glucose Tolerance, Insulin, and Lipid Profiles

In addition to reducing food intake and body weight, **GLP-1** is known to improve glucose handling [40,41]. To assess the efficacy of the four vectors on glucose handling, we measured blood glucose levels after an oral glucose challenge before and 3 weeks after dosing. At baseline, OGTT glucose excursions were comparable across all five groups (Figure 3A). At 3 weeks post-dose, both the **Ex4** and **Ex4-Fc** groups had lower fasting (time-0) glucose than vehicle (193 mg/dL, 144 mg/dL, 140 mg/dL for vehicle, Ex4, and Ex4-Fc, respectively) and lower glucose throughout the 120-minute challenge (Figure 3B). The reduction was particularly pronounced at 30 minutes in the **Ex4-Fc** group, which showed a substantially shortened peak (162 mg/dL at 30 minutes, versus 288 mg/dL in vehicle, Figure 3B). **GLP-1** and **Gly8** groups showed comparable glucose handling to the vehicle group (207 mg/dL, 217 mg/dL fasting and 298 mg/dL, 306 mg/dL at 30 minutes for **GLP-1** and **Gly8**, respectively). Comparing AUC relative to each group’s own baseline, week 3 AUC trended lower than baseline for all groups (Figure 3C). Though both the **Ex4** and **Ex4-Fc** groups showed larger reductions in AUC compared to vehicle, **GLP-1**, or **Gly8** groups (14.6%, 19.4%, 18.6%, 26.4%, 41.2% reductions in AUC for vehicle, **GLP-1, Gly8**, Ex4, **Ex4-Fc** groups respectively), only the **Ex4-Fc** group had a statistically significant (p < 0.01) improvement in glucose handling.

**Figure 3.**
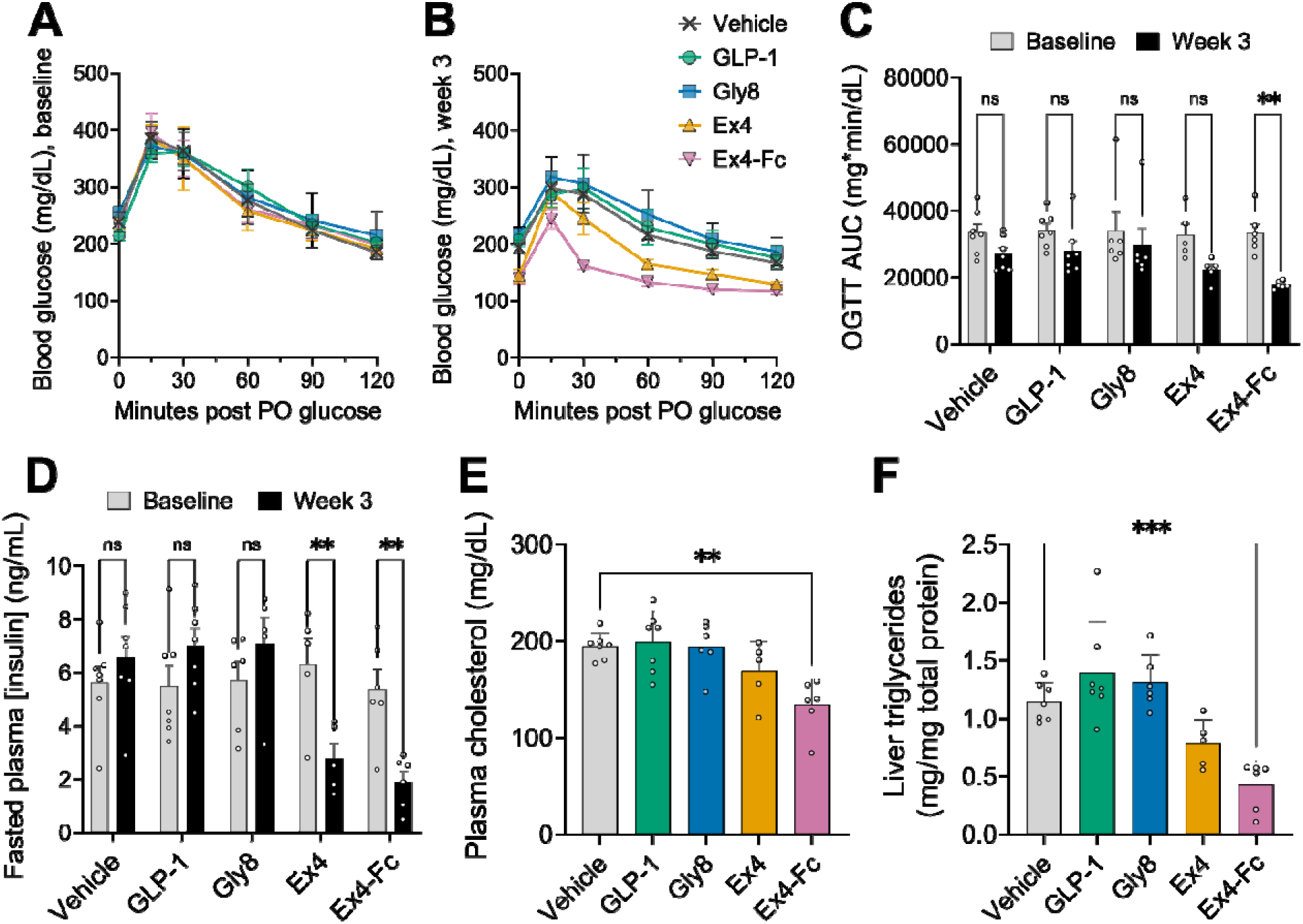
Glucose tolerance, insulin, and lipid profiles at 3 weeks post-dose. (A) OGTT blood glucose levels at baseline. (B) OGTT blood glucose levels at week 3. (C) Blood glucose AUC at baseline and week 3. (D) Fasted plasma insulin at baseline and week 3. (E) Terminal plasma cholesterol. (F) Liver triglycerides, normalized to total protein. Data were analyzed by two-way repeated-measures ANOVA with Šídák’s post hoc tests comparing baseline versus week 3 within each group (C, D) or one-way ANOVA with Dunnett’s post hoc tests versus vehicle (E, F). n = 5–7/group; ns, not significant; *p < 0.05; **p < 0.01; ***p < 0.001; ****p < 0.0001.

As would be expected from reduced fat mass, fasted plasma insulin was significantly reduced at 3 weeks relative to baseline in both the **Ex4** and **Ex4-Fc** groups (6.32 ng/mL vs. 2.77 ng/mL and 5.35 ng/mL vs. 1.89 ng/mL comparing baseline vs. week 3, **Ex4** and Ex4-Fc, respectively; p < 0.01 for each). In contrast, vehicle, **GLP-1**, and **Gly8** groups trended towards an increase in fasted plasma insulin on week 3 compared to baseline (Figure 3D). Additionally, both terminal plasma cholesterol (Figure 3E) and liver total triglycerides (Figure 3F) were significantly lower in the **Ex4-Fc** group relative to vehicle (134 mg/dL vs. 194 mg/dL and 0.43 mg/mg total protein vs 1.14 mg/mg total protein; p < 0.01 and p < 0.001, respectively). As with the other metabolic endpoints, the **Ex4** group also showed lower plasma cholesterol and liver triglycerides relative to the vehicle group. The effects seen in the **Ex4** group were smaller than those observed in the **Ex4-Fc** group and did not reach statistical significance.

### Transgene Biodistribution and GLP-1RA Exposure

We next investigated whether the lack of in vivo efficacy of the **GLP-1** and **Gly8** vectors could be explained by insufficient expression at the transcript level. To address this possibility, we quantified transgene transcripts in brain and liver by RT-dPCR across all five groups (Figure 4A). WPRE-targeted RT-dPCR transcript was detectable at high levels in both brain (4,542 ± 656.6 cp/ng RNA; pooled average ± SEM) and liver (2,468 ± 517.3 cp/ng RNA; pooled average ± SEM) across all four treatment groups. No WPRE transcripts were detected from vehicle-dosed brain or liver as expected. Brain transcript levels did not differ significantly across all pairwise comparisons (5,134 cp/ng RNA, 3,848 cp/ng RNA, 6,508 cp/ng RNA, 2,907 cp/ng RNA for **GLP-1, Gly8**, Ex4, Ex4-Fc, respectively). In liver, transgene transcript levels were the lowest in the **Ex4-Fc** group (161.0 cp/ng RNA). This was significantly lower than in the **GLP-1** group (3,675 cp/ng RNA; p < 0.05) and the **Ex4** group (2,041 cp/ng RNA; p < 0.05), ruling out failure of transgene transcription as the explanation for **GLP-1** vectors’ lack of in vivo efficacy. At the protein level, however, total **GLP-1** immunoreactivity (an assay that detects **GLP-1** (7-36) and the DPP-IV cleavage product (9-36) but not Exendin-4) in plasma (Figure 4B) and brain lysate (Figure 4C) was similar across all five groups, including vehicle (57.1 pM, 43.7 pM, 58.6 pM, 41.2 pM, 62.2 pM for plasma [**GLP-1**] and 3.17 fmol/mg, 3.45 fmol/mg, 4.41 fmol/mg, 3.32 fmol/mg, 2.96 fmol/mg for brain lysate [**GLP-1**] in vehicle, **GLP-1, Gly8**, Ex4, **Ex4-Fc** animals, respectively). We therefore conclude that, at the modest dose tested (5×10^12^ vg/kg), neither the **GLP-1** nor the **Gly8** vector raised total **GLP-1** immunoreactivity above endogenous levels in plasma or brain, despite high transcript levels in both tissues.

**Figure 4.**
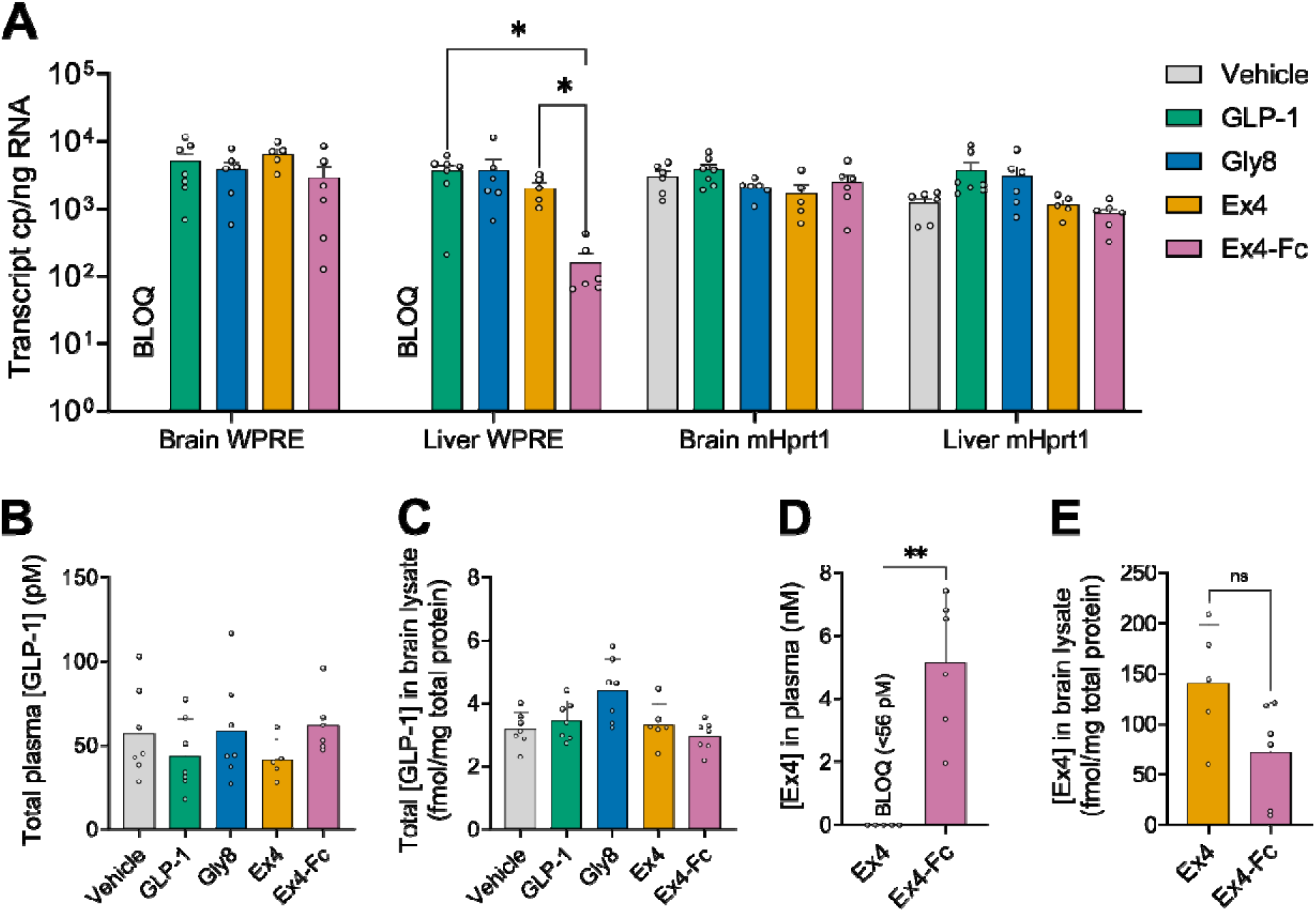
Transgene biodistribution and GLP-1RA exposure at study termination. (A) WPRE3-targeted RT-dPCR transcript in brain and liver, alongside mHprt1 reference transcript. (B) Total plasma [**GLP-1**], determined by ELISA. (C) Total **GLP-1** in brain lysate, by ELISA. (D) Exendin-4 in plasma of **Ex4** and **Ex4-Fc** groups; samples below the limit of quantification (BLOQ) are plotted at the assay lower limit of quantification (0.056 nM). (E) Exendin-4 in brain lysate of **Ex4** and **Ex4-Fc** groups. Data were analyzed by two-way ANOVA with Šídák’s post hoc tests comparing WPRE transcript copy number within each tissue group (A) or one-way ANOVA with Dunnett’s post hoc tests versus vehicle (B, C), or Student’s t-test (D, E). n = 5–7/group; ns, not significant; *p < 0.05; **p < 0.01.

Separately, we investigated whether **Ex4** and **Ex4-Fc** differed in central versus peripheral exposure that could account for the differences observed in the functional measurements. Despite having the lowest WPRE transcript levels in the liver out of the four treatment groups (Figure 4A), the **Ex4-Fc** group had higher levels (5.14 nM) of detectable protein in terminal plasma than the **Ex4** group (p < 0.01; Figure 4D). All plasma samples measured from the **Ex4** group had peptide levels below the assay’s limit of quantification (<0.056 nM). In brain lysates, by contrast, the **Ex4** group had higher protein levels than observed in the **Ex4-Fc** group (140.7 fmol/mg vs. 72.0 fmol/mg; Figure 4E), though this difference was not significant.

## Discussion

In the present work, we compared the in vitro potency, in vivo efficacy, and tissue-level distribution of four AAV-vectorized GLP-1RAswith varying degrees of engineering of the peptide sequence. Two constructs, **Ex4** and Ex4-Fc, resulted in weight loss, fat-mass reduction, and reduced fasting insulin observed up to 3 weeks after a single dose of 5×10^12^ vg/kg. This dose, along with capsid and promoter, was held constant across all four constructs so that differences in outcome could be attributed to the vectorized peptide sequence itself rather than to differences in vector exposure. The best performing construct, Ex4-Fc, resulted in 22.6% body weight reduction at day 21. These data are consistent with literature data which show that repeated injections of 100 nmol/kg/day semaglutide in DIO mice can achieve 22% body weight loss at 3 weeks [42]. Native **GLP-1** and **Gly8** vectors resulted in no detectable metabolic phenotype, despite confirmed secretion of potent **GLP-1**R agonists in vitro and transcript levels comparable to **Ex4** and **Ex4-Fc** in brain and liver. Comparing **GLP-1** protein levels in plasma and brain across groups suggests that the lack of efficacy observed with **GLP-1** and **Gly8** can most likely be attributed to either poor stability in vivo or poor translation efficiency.

The comparison of vectorized **GLP-1, Gly8**, and **Ex4** was intended to test whether constitutive local production could overcome the peptidase liability of the native sequence in vivo. Three possible drivers of the poor performance of **GLP-1** and **Gly8** relative to **Ex4** and **Ex4-Fc** are considered below: DPP-IV liability, secretion efficiency, and cleavage by other peptidases such as NEP. First, the in vitro comparison of **GLP-1** and **Gly8** provides some evidence against DPP-IV liability as the bottleneck in our study. **GLP-1** and **Gly8** had similarly low functional GLP-1RA levels in vitro despite the known DPP-IV resistance conferred by the A8G substitution. Additionally, the ELISA used to measure total secreted GLP-1RA also detects the DPP-IV cleavage product **GLP-1**(9-36), but the total immunoreactive and **GLP-1**R active GLP-1RA levels were orders of magnitude lower for both **GLP-1** and **Gly8** than for **Ex4** or Ex4-Fc. Second, secretion efficiency is also unlikely to be the primary driver: all four constructs used the same signal peptide and the same N-terminal residue — the dipeptide processed by signal peptidase was identical across constructs. Lysate concentrations closely tracked media concentrations for every construct, with [media]/[lysate] ratios differing by at most 3-fold between **GLP-1** and Ex4-Fc, further downplaying secretion efficiency as the main driver of the drastic differences in efficacy across constructs. Third, neutral endopeptidase 24.11 (NEP) could account for at least some of the differences observed between the constructs: NEP can cleave both **GLP-1** and **Gly8** at multiple internal sites but is comparatively poor at cleaving **Ex4** [10,13]. This aligns well with our observation that both **Ex4** and **Ex4-Fc** performed well in terms of in vitro potency and in vivo efficacy and both **GLP-1** and **Gly8** did not. It is also possible that other peptidases, such as carboxypeptidase A2, may play an oversized role when dealing with local GLP-1RA expression outside of prohormone context.

An alternate explanation for the discrepancies across in vivo efficacies could be that **GLP-1** and **Gly8** are produced far less efficiently at the level of early secretory-pathway processing, independent of downstream peptidase liability. Small, unstructured peptide hormones like **GLP-1** are typically synthesized as part of a larger prohormone and matured only late in the secretory pathway. Expressing the mature 30-residue peptide directly, without that prohormone context, would leave its hydrophobic residues exposed to ER-associated degradation (ERAD) pathway or aggregation [43]. In contrast, Exendin-4 has a C-terminal Trp-cage that provides more structure and improved solubility [44]. Consistent with our observation, other groups have previously reported that expressing **GLP-1** without further engineering leads to poor secretion [45,46]. C-terminal fusions of well-folded proteins such as nanobodies or Fc domains have been reported to generally improve the production of vectorized peptides, which normally produce poorly [47,48]. The superiority we observed of **Ex4-Fc** over **Ex4** in both in vitro secretion and in vivo efficacy demonstrates that the overall production of vectorized GLP-1RAsin transduced cells is a relevant parameter that can be optimized in vitro. A prior study using a native **GLP-1** AAV saw minor improvements in glucose handling but not weight loss in db/db mice at a much higher dose [49], suggesting that the therapeutic effect is theoretically achievable with a near-endogenous sequence given high enough expression levels. However, this previous **GLP-1** AAV also included a furin-cleavable hemagglutinin fusion that greatly improved secretion compared to the naked **GLP-1** peptide sequence in vitro.

**Ex4** and **Ex4-Fc** demonstrated that a vectorized GLP-1RA can be efficacious: a single intravenous dose produced sustained weight loss, fat-mass reduction, and improved metabolic function for the full 3-week study period. The magnitude of peak weight loss achieved — 18.9% and 22.6% of starting body weight for **Ex4** and Ex4-Fc, respectively — is comparable to that reported for the drug semaglutide in diet-induced obese rodents [42], and the overall efficacy of **Ex4** is consistent with prior reports of AAV-delivered Exendin-4 in rodent models of diet-induced obesity and type 2 diabetes [50]. These results support the central premise of this study: a single administration of a sufficiently well-engineered GLP-1RA transgene can recapitulate the metabolic benefit of chronic pharmacologic GLP-1R agonism. **Ex4** and **Ex4-Fc** showed a notable mismatch between transcript and protein. Despite significantly higher liver transcript levels in the **Ex4** group, plasma exposure was reversed: every **Ex4** plasma sample fell below the assay’s lower limit of quantification (<0.056 nM), at least 90-fold below average **Ex4-Fc** plasma levels. This is consistent with the IgG Fc domain’s established role in extending systemic half-life, since Fc fusion raises the protein’s size above the ∼60 kDa renal filtration cutoff and enables FcRn-mediated recycling [51,52]. Taken at face value, the plasma exposure data raise a separate question: how do **Ex4** and **Ex4-Fc** produce comparable hypophagia in week 1 when **Ex4** plasma levels are below the limit of quantification at the end of study? Hypophagia downstream of **GLP-1**R agonism is thought to be driven centrally by multiple neuronal subpopulations in the hypothalamus, hindbrain, and the lateral septum [17,18,53]. While some of these target areas, such as the area postrema, have some direct access to GLP-1RAsin systemic circulation via fenestrated capillaries [54], others, such as the lateral septum, are restricted to GLP-1RAsin the CNS proper. Central and peripheral exposure may therefore drive hypophagia, consistent with **Ex4** and **Ex4-Fc** producing weight loss of similar magnitude despite their divergent exposure profiles. Downstream of hypophagia and weight loss, glucose handling, insulin, and lipids would be expected to move as a secondary consequence of reduced feeding and fat loss, regardless of circulating GLP-1RA levels. Ex4-Fc’s more robust peripheral phenotype — OGTT improvement, cholesterol and triglyceride reduction — may additionally reflect direct peripheral **GLP-1**R engagement in pancreas, liver, and vagal afferents [19,20], on top of the central hypophagia pathways it shares with Ex4.

Both the **Ex4** and **Ex4-Fc** groups showed a gradual loss of hypophagia between weeks 2 and 3, which is consistent with a gradual increase in food intake over the course of chronic GLP-1RA dosing reported previously with both Exendin-4 and semaglutide [55,56]. This pattern is also consistent with homeostatic defense of a body weight set point, in which compensatory reductions in energy expenditure and/or increases in appetite erode drug-induced hypophagia over time, independent of receptor-level pharmacology [57,58]. What is surprising is that weight regain occurred in every animal dosed with **Ex4** AAV despite high viral transcript levels in brain and liver across all treatment groups. Promoter silencing is a known phenomenon with CMV promoter used in this study but unlikely to explain the weight regain in the **Ex4** group. AAV-driven transgene mRNA and protein in mouse brain rise over the first three weeks before peaking [59] whereas CMV methylation-dependent silencing shows earliest reduction of expression after four weeks [60]. Consistent with this, WPRE transcript remained readily detectable in brain and liver at week 3 for all groups. **GLP-1**R is a class B GPCR, and continuous, non-pulsatile agonism of this receptor family is known to drive ligand-induced internalization and homologous desensitization [61,62,63,64]. One possibility is that non-pulsatile **GLP-1**R agonism without a drug holiday reduces the available receptor pool and right-shifts the dose-response curve. Under this scenario, only the **Ex4-Fc** group, whose plasma levels were 90-fold higher than the **Ex4** group at study end, would retain exposure sufficient for continued efficacy.

Anti-drug antibody (ADA) formation is a known liability of Exendin-4 and offers another plausible explanation for the loss of efficacy in **Ex4** AAV-dosed animals. This explanation sits uneasily, however, with the observation that **Ex4-Fc** (which carries the larger, more antigenic human IgG4 Fc domain) showed no comparable weight regain over the same interval. The rate of regain is also difficult to reconcile with reported ADA incidence: every animal dosed with **Ex4** AAV in the present study regained weight between weeks 2 and 3, whereas ADA formation against Exendin-4 in mice is reported at a substantially lower frequency (≤10%, per the FDA pharmacology/toxicology review of exenatide) [65].

ADA formation is a meaningful risk for any AAV-delivered therapeutic, since dosing cannot be reduced or withdrawn the way it can for a conventional biologic once an immune response develops. There is, however, growing evidence that AAV-mediated transgene expression from hepatocytes can itself drive a tolerizing response toward transgene-derived proteins through a CD25+ regulatory T-cell-dependent mechanism, an approach already explored for hepatic AAV gene immunotherapy in food allergy [66]. With the right combination of capsid, a non-constitutive promoter (e.g., the glucose-sensitive insulin promoter used in Fractyl Health’s platform [67]), and transgene design, it may be possible to engineer an AAV-GLP-1RA therapeutic with both durable efficacy and low ADA risk.

The results of the present study are an encouraging proof of concept: a single intravenous dose of **Ex4-Fc** AAV produced substantial, durable weight loss and metabolic improvement in obese mice, with no overt signs of ill health over the 3-week study period. Two principles from this work are likely to generalize to vectorizing secreted therapeutics more broadly.

First, amino-acid-level sequence engineering remains important even under constitutive local production. Optimizing the production and secretion efficiency of the therapeutic peptide is a critical early engineering step, one that can be screened in vitro and translates well to in vivo efficacy. Second, the in vivo biodistribution of a vectorized therapeutic is not a property of the capsid and promoter alone but rather the convolution of capsid tropism, promoter activity, and protein/peptide stability. This makes biodistribution difficult to predict from any single component and can produce sharply different patterns at the transcript versus protein level. When leveraged deliberately, tuning capsid, promoter, and payload together could allow for high CNS and low plasma exposure: a pattern that is difficult to achieve by delivering peptide directly but was observed for the **Ex4** group.

## Funding

This work was funded by Neurocrine Biosciences, Inc. Neurocrine Biosciences, Inc. was involved in the analysis of the data and the preparation of the manuscript.

## Conflict of Interest

All authors are employees of Neurocrine Biosciences, Inc. and may own stock or stock options in the company.

## Data Availability Statement

The data supporting the findings of this study are proprietary to Neurocrine Biosciences, Inc. and are not publicly available due to company confidentiality policies.

## Author Contributions

DL: Conceptualization, Methodology, Investigation, Formal analysis, Visualization, Writing – original draft. JO, KB, AM: Conceptualization, Supervision. GN, TB: Conceptualization, Methodology, Resources, Supervision. ET: Methodology, Supervision. JB, LE, BB: Investigation. JS, VL: Supervision. DW: Conceptualization, Resources, Supervision. All authors: Writing – review & editing.

## Abbreviations

AAV: adeno-associated virus
ADA: anti-drug antibody
ANOVA: analysis of variance
AUC: area under the curve
bGH polyA: bovine growth hormone polyadenylation signal
BLOQ: below the limit of quantification
CMV: cytomegalovirus
CNS: central nervous system
DIO: diet-induced obese
DPP-IV: dipeptidyl peptidase-IV
ELISA: enzyme-linked immunosorbent assay
ERAD: ER-associated degradation
Ex4: Exendin-4
Ex4-Fc: Exendin-4–human IgG4 Fc fusion
FcRn: neonatal Fc receptor
GIP: glucose-dependent insulinotropic polypeptide
GLP-1: glucagon-like peptide-1
GLP-1R: GLP-1 receptor
GLP-1RA: GLP-1 receptor agonist
GPCR: G protein-coupled receptor
HFD: high-fat diet
ITR: inverted terminal repeat
MANOVA: multivariate analysis of variance
NEP: neutral endopeptidase 24.11
OGTT: oral glucose tolerance test
RT-dPCR: reverse-transcription digital PCR
scAAV: self-complementary AAV
TR-FRET: time-resolved fluorescence resonance energy transfer
vg: vector genomes
WPRE: woodchuck hepatitis virus posttranscriptional regulatory element.

---

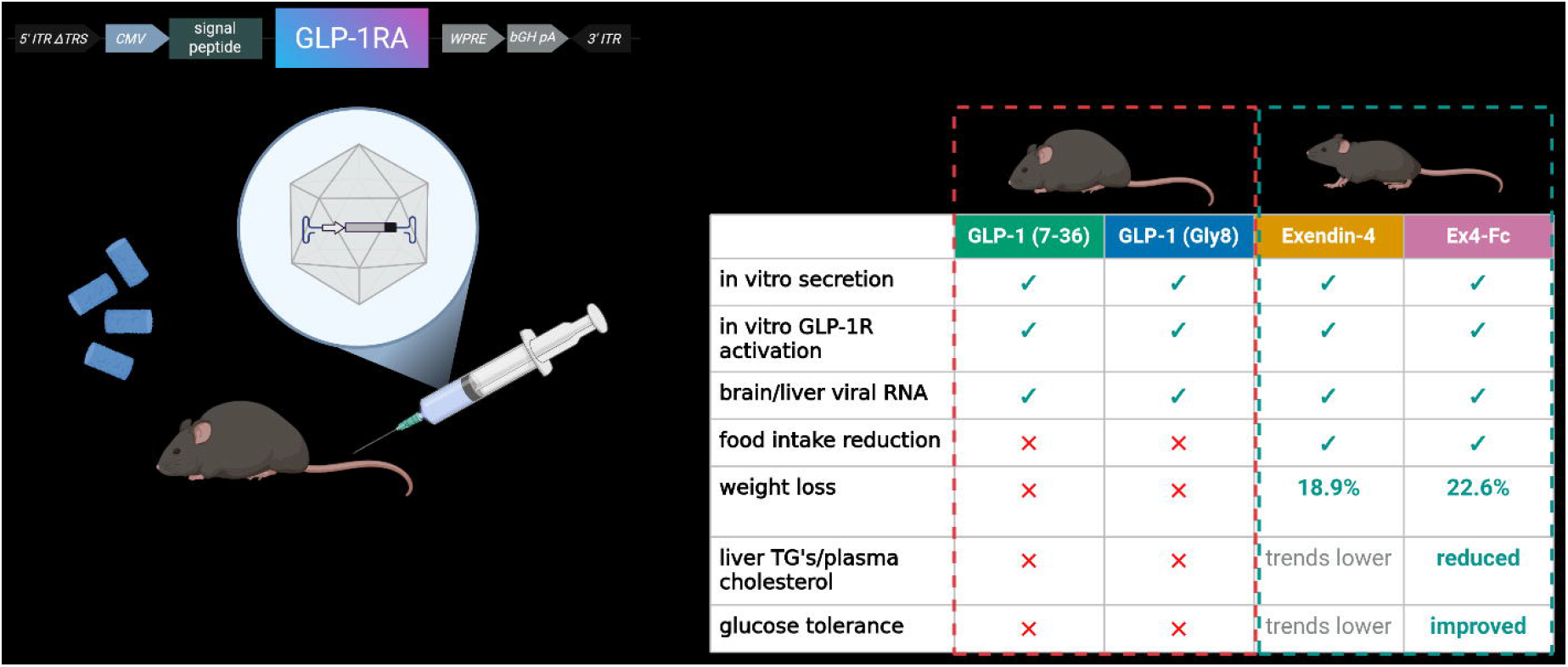

## Notes

### Competing Interest Statement

D.L., K.B., G.N., A.M., T.B., E.T., J.B., J.S., V.L., L.E., B.B., J.O., and D.W. are employees of Neurocrine Biosciences, Inc. and may own stock in the company.

